# Bacterial profiles of the oral, vaginal, and rectal mucosa and colostrum of periparturient sows

**DOI:** 10.1101/2024.03.27.586787

**Authors:** Virpi Piirainen, Emilia König, Aleksi Husso, Mari Heinonen, Antti Iivanainen, Tiina Pessa-Morikawa, Mikael Niku

**Affiliations:** Department of Production Animal Medicine, Faculty of Veterinary Medicine, University of Helsinki, Helsinki, Finland; Department of Veterinary Biosciences, Faculty of Veterinary Medicine, University of Helsinki, Helsinki, Finland; Research Centre for Animal Welfare, Department of Production Animal Medicine, University of Helsinki, Helsinki, Finland

## Abstract

The commensal microbiota influences the health, feeding efficiency, and reproductive performance of sows. The microbiota composition in the alimentary and genitourinary tracts and in colostrum/milk during pregnancy and lactation also impacts the developing microbiota and immune system of the piglets and subsequently their growth and health. Knowledge of the microbial compositions is important for evaluation of these effects and for discovering ways to improve the health and productivity of the sows.

Oral, vaginal, and rectal mucosa were sampled from 32 sows of variable parity in late pregnancy on four commercial piglet-producing farms in Finland. Colostrum samples were taken within 6 hours of delivery of the first piglet. Microbial compositions were analyzed by 16S rRNA gene amplicon sequencing. Moderate differences in diversity and composition were observed between farms. The most abundant genera of the oral microbiota were *Rothia*, *Moraxella*, and *Streptococcus.* The rectal microbiota was dominated by *Clostridium sensu stricto 1. Streptococcus* was the most abundant genus in the vagina and colostrum. Differences in relative abundances of genera were detected between primiparous and multiparous sows. Some of these differences were detected across all the farms; in the multiparous sows, the relative abundances of *Fusobacterium* and *Neisseria* were lower in oral samples. *Clostridium sensu stricto 1*, *Romboutsia*, and *Lachnospiraceae_UCG_007* were higher and *Prevotella* lower in rectal samples, and *Streptococcus* higher in colostrum samples. In vaginal samples, approximately half of the multiparous sows had significantly higher relative abundances of the genera *Fusobacterium* and *Streptococcus* than the primiparous sows. Among the differentially abundant taxa, *F. necrophorum* and *F. nucleatum* were identified in oral samples, *Fusobacterium gastrosuis* and *Fusobacterium necrophorum* in vaginal samples, and *Streptococcus dysgalactiae* in colostrum samples. Most of the differences were due to unidentified species within the respective genus.

This study provides a comprehensive overview of the mucosal and colostrum microbiota of periparturient sows during normal production conditions on Finnish commercial farms, including potentially interesting differences in the relative abundances of several genera at different mucosal sites between sows of low and high parity.

## Background

The mucosal surfaces, skin, feces, and glandular excretions of the mammalian body are all inhabited by distinct communities of resident microbes. Once established early in life, these microbiotas are relatively stable [1-3]. Commensal microbes provide health and wellbeing benefits to the host by improving maintenance of the epithelial barrier [4] and providing protection against pathogens [5]. They also influence host immune development and immune functions [6-9] and feed utilization [10]. Furthermore, commensal microbes have effects on neural development and function and may even impact host mood and behavior [11, 12].

Due to the potential benefits of the microbiota on the health and performance of the host, there has been a growing interest in the composition of the commensal microbiota of production animals, including pigs. These studies have mostly focused on growing pigs [13, 14], but the commensal microbiota of pregnant sows has also gained attention to an increasing extent [15-24]. Physiological factors, including phase of the reproduction cycle [16-19] and parity [21-23], can modulate the microbial compositions of the adult animals.

Environmental factors, such as diet, stress, and exposure to antibiotics [13, 25-27], can also have impacts. The fecal, vaginal, and colostrum (or milk) microbiotas of the sow in turn influence the developing microbiota and immune system of the offspring during pregnancy and lactation [6, 22, 24, 28, 29]. This has long-term effects on the health and growth of the piglets [6, 30]. Thus, knowledge on sow microbiota composition is valuable in evaluating its potential benefits and disadvantages to the offspring.

In this study, we explored the bacterial profiles of the oral, vaginal, and rectal mucosa and colostrum of 32 periparturient sows using 16S RNA gene amplicon sequencing. The study population included both primiparous and multiparous sows from four commercial Finnish piglet-producing farms. The bacterial profiles were compared between farms, parity groups, and individuals.

## Results

### Overview of sequencing data

16S rRNA gene amplicon sequencing of sow oral, vaginal, and rectal mucosa and colostrum samples (n = 126) resulted in 6 749 409 high-quality reads, with an average of 53 567 (± SD 12 960) reads per sample. The reads were mapped to 5379 ASVs. Details of the sequencing data for each sample type are shown in Supplementary file 1, Supplementary table 1.

### Microbiota diversity in different mucosal sites

The alpha diversity indices of the four sample types are shown in Fig 1a. Each sample type clustered separately in PCoA (Fig 1b). In PCoA, the oral samples were completely separated from the other sample types on the PCo1 axis. Vaginal samples were separated from the others on the PCo2 axis, with some overlap with the rectal cluster (Fig 1b). The colostrum samples clustered closest to but separate from the rectal samples and separate from the oral and vaginal samples (Fig 1b).

**Figure 1.**
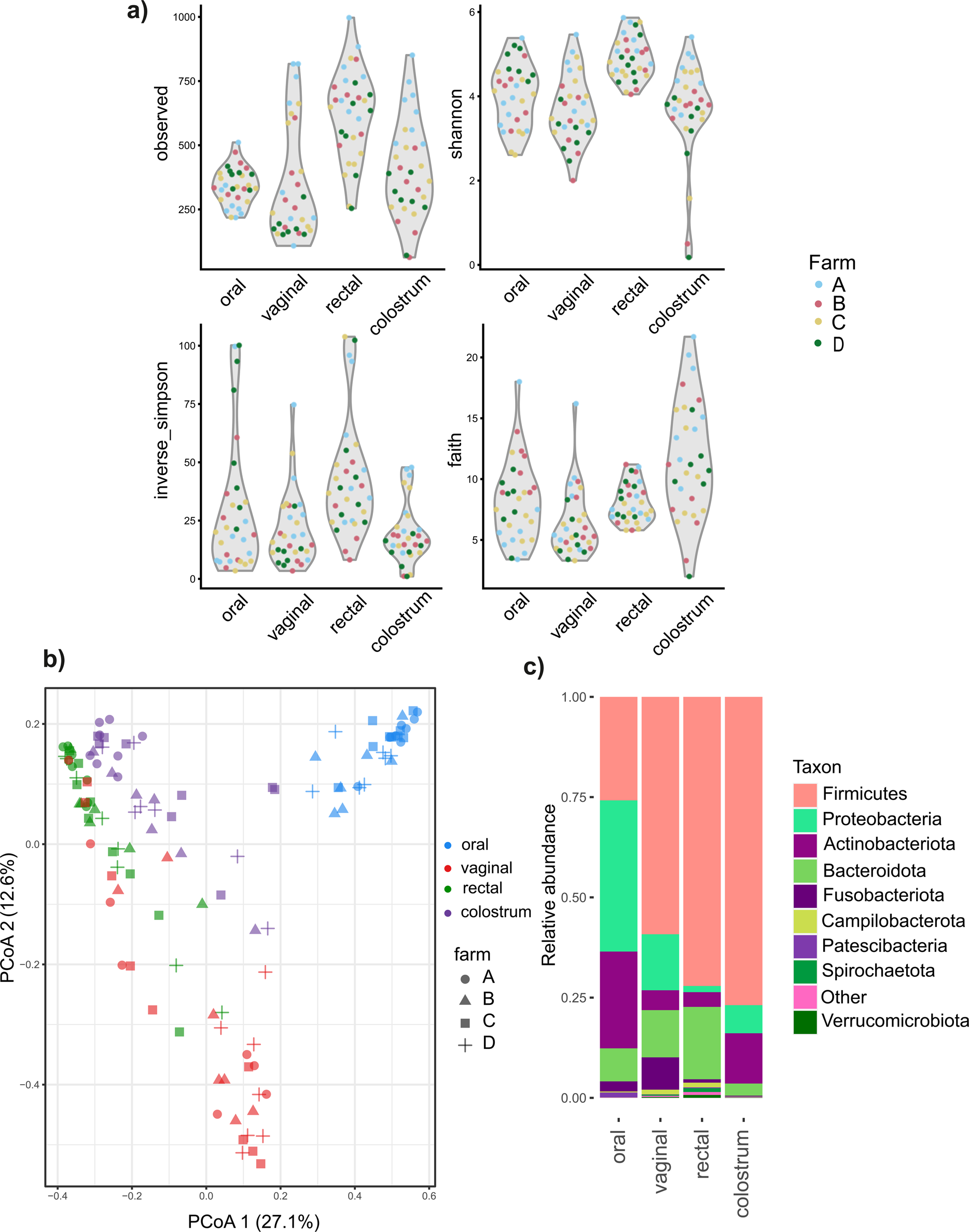
Overview of the oral, vaginal, rectal, and colostrum microbiota in 32 pregnant sows from four Finnish commercial farms, A-D. a) Alpha diversity, b) PCoA of Bray Curtis dissimilarities, c) relative abundances of major phyla.

Nine bacterial phyla were identified in the oral samples, 11 in vaginal, 15 in rectal, and 11 in colostrum samples. Firmicutes, Bacteroidota, Proteobacteria, and Actinobacteriota were the dominant phyla in all sample types. In the oral samples, Proteobacteria were the most abundant phylum (38%), followed by Firmicutes (26%) and Actinobacteriota (24%). In the vaginal samples, Firmicutes (59%) were the most abundant phylum, followed by Proteobacteria (14%) and Bacteroidota (12%). In rectal samples, Firmicutes (72%) were the dominant phylum, followed by Bacteroidota (18%) and Actinobacteriota (3.7%). Firmicutes (77%) were also dominant in the colostrum samples, followed by Actinobacteriota (13%) and Proteobacteria (7.0%). The overall relative abundances (RAs) of the main phyla in each sample type are shown in Fig 1c.

### Sow oral microbiota

The median alpha diversity of farm D samples was higher than that of the others (Fig 2a), but the difference was significant only between farms D and C. In PCoA, the samples from different farms overlapped, but a partial separation of farm D samples from the others on PCo1 and of farm B on PCo2 could be observed (Fig 2b).

**Figure 2.**
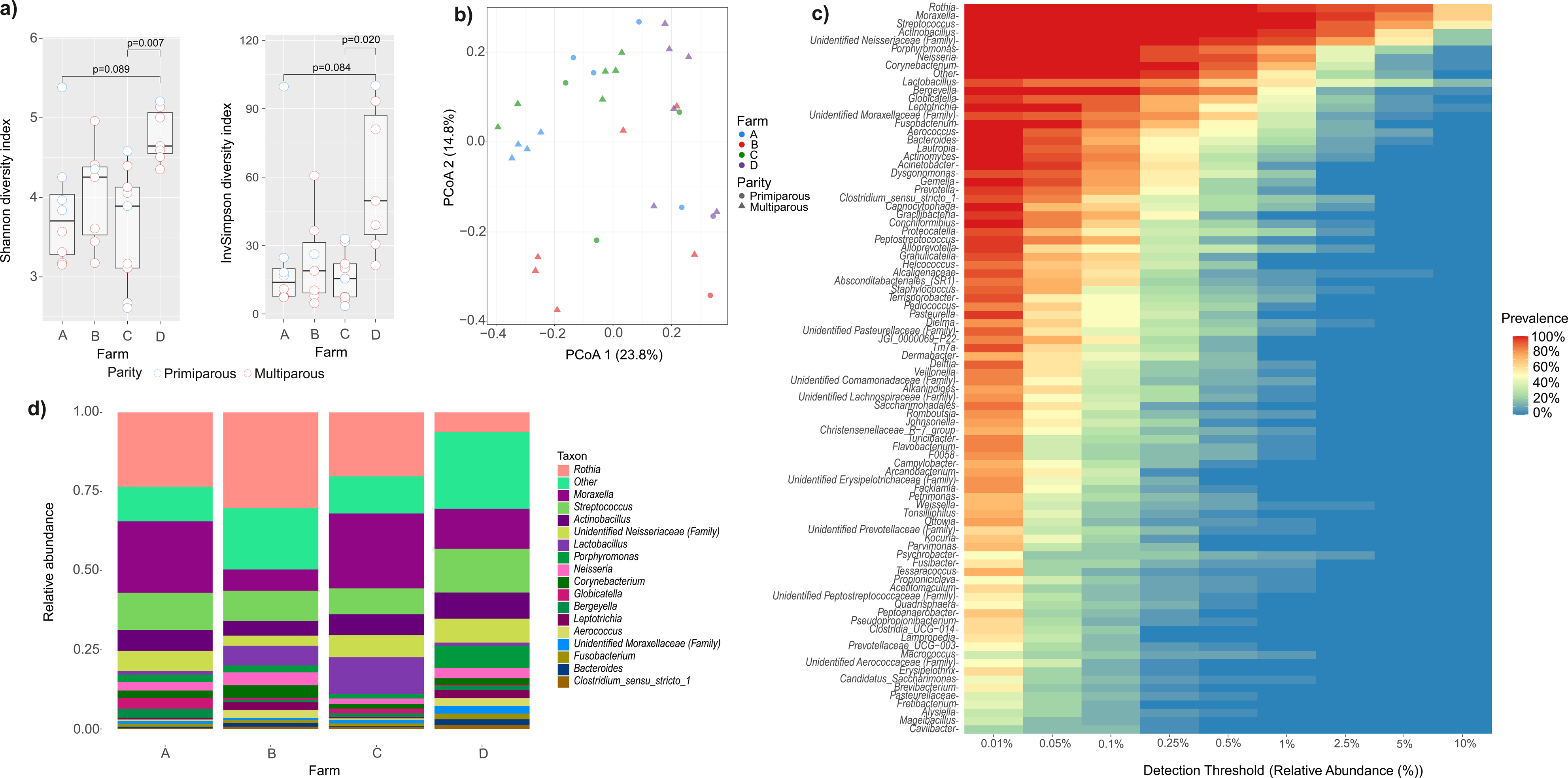
Composition of the oral microbiota of pregnant sows from four Finnish commercial farms. a) Alpha diversity indices Shannon and Inverse Simpson, b) principal coordinates analysis based on Bray-Curtis dissimilarities, c) Core heatmap, d) Average relative abundances of the main genera in farms A-D.

In total, 113 genera were identified in the oral samples. The microbiota composition at the genus level is presented in Fig 2c. Twenty-one genera had 100% prevalence and together constituted 79% of the total abundance calculated from the RA data. The most abundant of these genera were *Rothia*, *Moraxella*, *Streptococcus*, *Actinobacillus*, an unidentified genus of the *Neisseriaceae* family, *Porphyromonas*, *Neisseria* and *Corynebacterium*, all present with RA >0.1% (Fig 2c). There were some apparent differences in the genus-level compositions between the farms with farm A and D sows having higher RA of *Moraxella* and farm D lower RA of *Rothia* than sows from the other farms (Fig 2d). Further, sows from farms A and D had lower RAs of *Lactobacillus* than those of farms B and C (Fig 2d).

### Sow vaginal microbiota

There were modest differences in the microbial diversities between the sows from the different farms, with farm A and C samples having higher alpha diversity than the others (Fig 3a). Only the difference between farm C and D was significant in Shannon. No differences were significant between the farms in Inverse Simpson’s. The farms were not separated in PCoA, but three primiparous sows from farm A clustered separately on PCoA 1 (Fig 3b).

**Figure 3.**
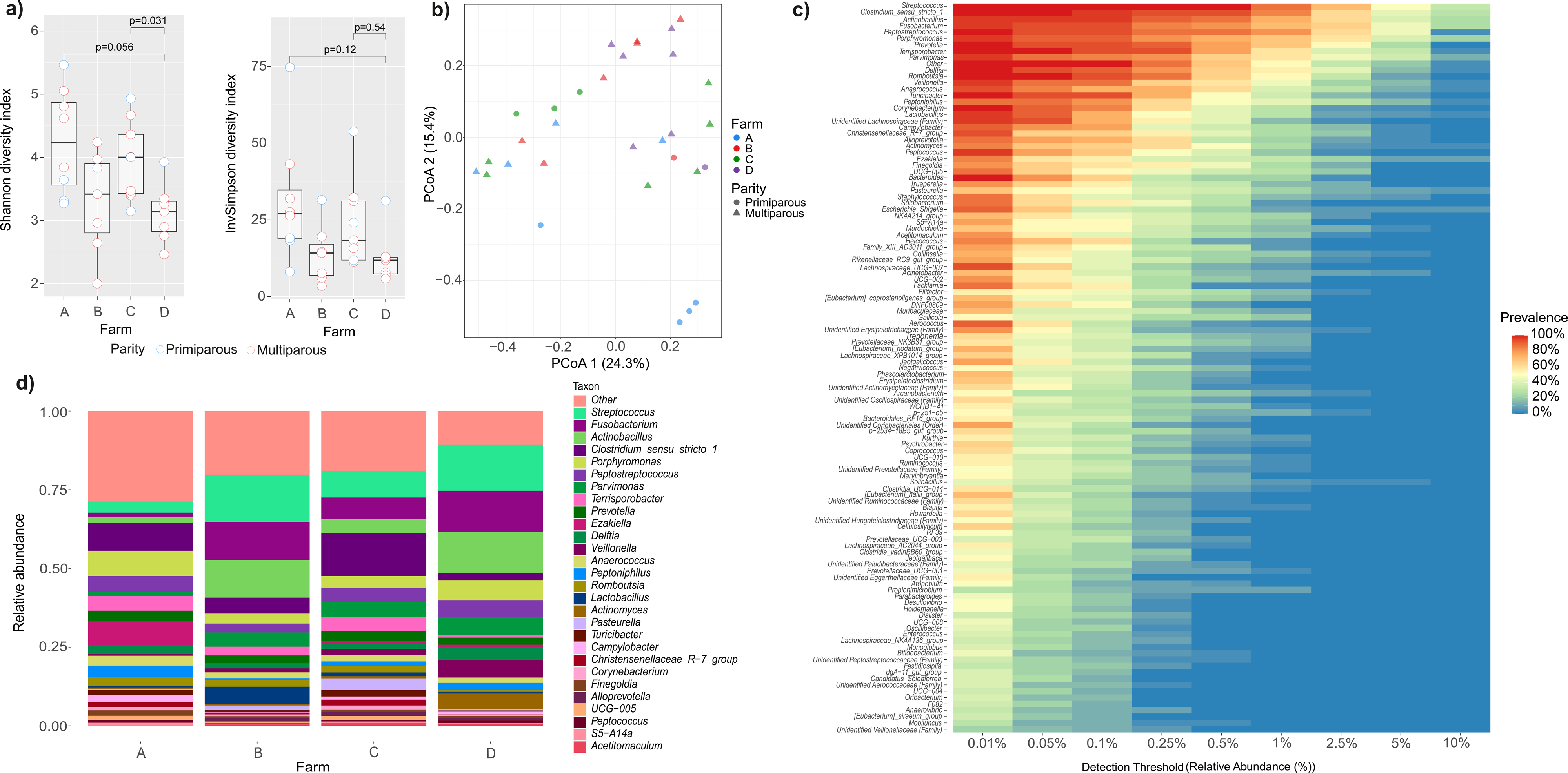
Composition of the vaginal microbiota of pregnant sows from four Finnish commercial farms. a) Alpha diversity indices Shannon and Inverse Simpson, b) Principal coordinates analysis based on Bray-Curtis dissimilarities, c) Core heatmap, d) Average relative abundances of the main genera in farms A-D.

In vaginal samples, 129 genera were identified. Altogether 13 of these were present in all samples constituting 42% of the total microbiota. *Streptococcus*, *Clostridium sensu stricto 1*, *Peptostreptococcus*, *Parvimonas*, *Terrisporobacter*, *Prevotella*, *Delftia*, *Romboutsia*, *Lactobacillus*, and *Turicibacter* were the most abundant of these genera (Fig 3c).

*Actinobacillus* and *Fusobacterium* had also high relative abundances but remained undetected in one sample. Farm A samples had lower RAs of *Streptococcus*, *Fusobacterium*, and *Actinobacillus* than the others, but higher RAs of *Porphyromonas*, *Ezakiella* and *Peptoniphilus*. Farms B and D had higher RAs of *Streptococcus*, *Fusobacterium*, and *Actinobacillus* than farms A and C, and higher RA of *Lactobacillus* compared to other farms (Fig 3d).

### Sow rectal microbiota

Although the rectal microbiota in farm A was more diverse than that of the other farms (Fig 4a), the differences in the diversity indices were not significant. Farm A and C and partially also farm B samples separated from each other in PCoA, with farm D samples spread among all the other farm groups (Fig 4b).

**Figure 4.**
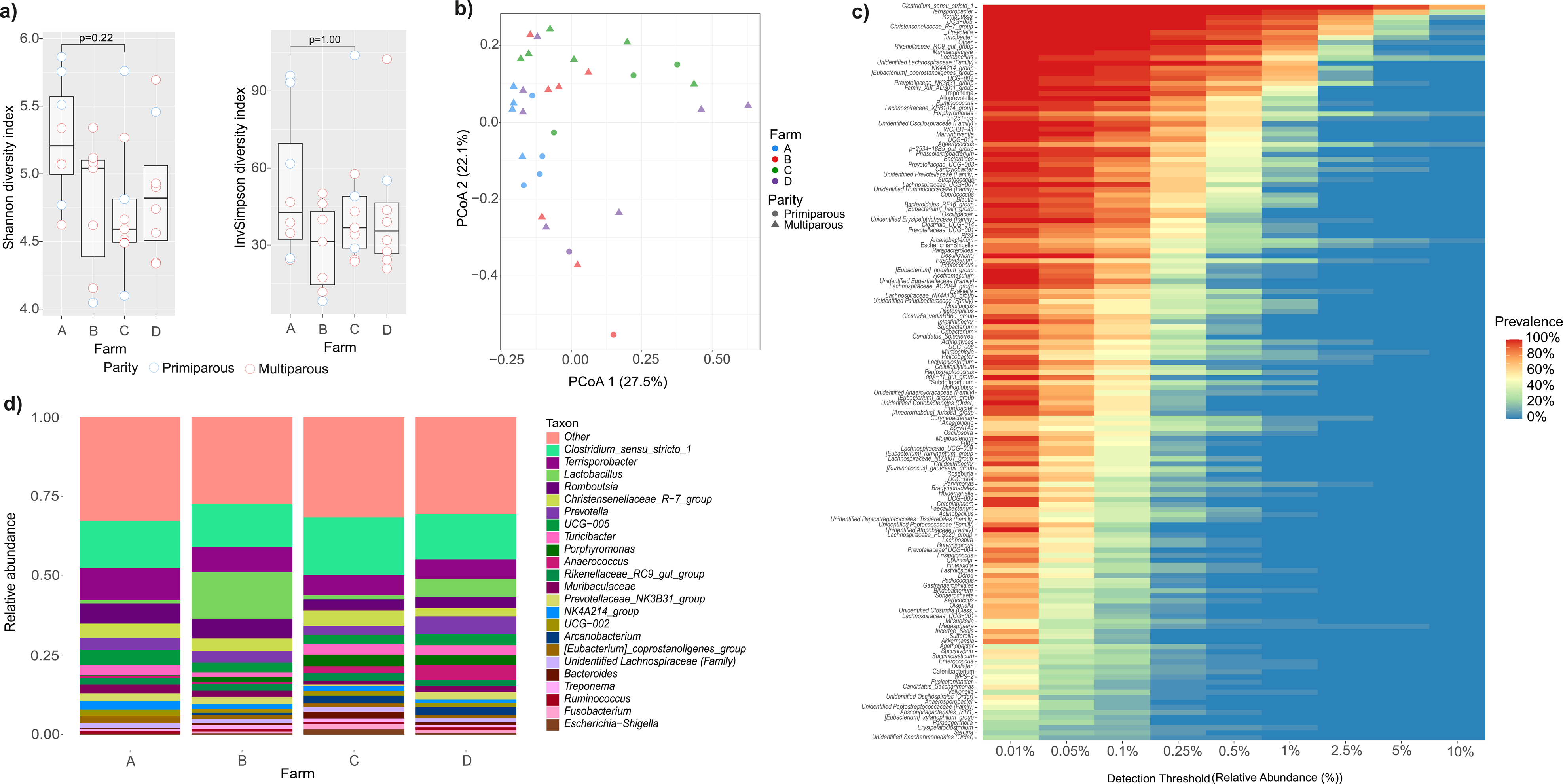
Composition of the rectal microbiota of pregnant sows from four Finnish commercial farms. a) Alpha diversity indices Shannon and Inverse Simpson, b) Principal coordinates analysis based on Bray-Curtis dissimilarities, c) Core heatmap, d) Average relative abundances of the main genera in farms A-D.

Altogether 161 genera were identified in the sow rectal samples. Forty-one of these had 100% prevalence and accounted for 71% of the total abundance. The most abundant of these were *Clostridium sensu stricto 1*, *Terrisporobacter*, *Lactobacilllus*, *Romboutsia*, UCG-005, *Christensenellaceae*_R−7_group, *Prevotella*, *Turicibacter,* and *Rikenellaceae*_RC9_gut_group. Farm B samples had higher abundances of *Lactobacillus* and lower of *Turicibacter* than samples from the other farms (Fig 4d). The RA of *Lactobacillus* in samples from farm D was also higher than that of the samples from farms A and C. Farm A and B samples had more *Romboutsia* than the others (Fig 4d).

### Sow colostrum microbiota

Although colostrum samples from farm A showed higher alpha diversity than others, the difference from the other farms was not significant (Fig 5a). In PCoA, most of the samples formed a cluster on PCo1, with three outliers belonging to two different farms clustering separately in the upper right corner (Fig 5b).

**Figure 5.**
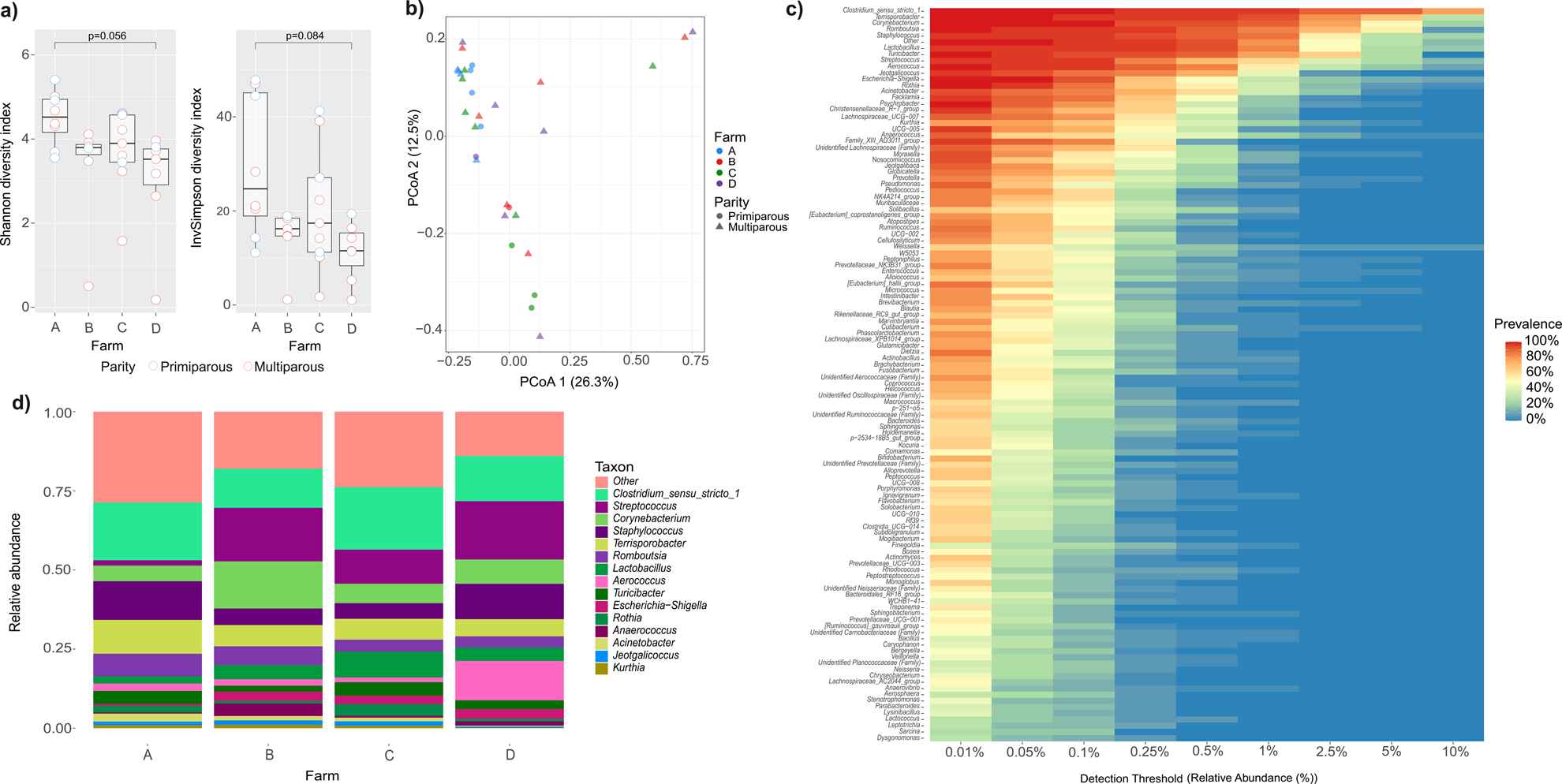
Composition of the colostrum microbiota of pregnant sows from four Finnish commercial farms. a) Alpha diversity indices Shannon and Inverse Simpson, b) Principal coordinates analysis based on Bray-Curtis dissimilarities, c) Core heatmap, d) Average relative abundances of the main genera in farms A-D.

Among the 159 genera identified, 13 were present in all samples and accounted for 64% of the total abundance. *Clostridium sensu stricto 1*, *Terrisporobacter*, *Corynebacterium*, *Romboutsia*, and *Staphylococcus* were the most abundant genera (Fig 5c). Samples from farm A had lower abundance of *Streptococcus* and higher abundance of *Staphylococcus* and *Terrisporobacter* than other farms. Samples from farm B had higher RA of *Corynebacterium* and *Anaerococcus*, and lower RA of *Clostridium* than the others (Fig 5d). Farm C had more *Rothia* than other farms, and farm D more *Aerococcus* (Fig 5d).

### Parity effects on microbial compositions

The alpha diversities between primiparous and multiparous sows were not significantly different, and the groups overlapped in PCoA (Supplementary file 2, Fig 1). However, we detected differences in the RAs of individual genera in each of the sample types. RAs of the main genera in all samples sorted by parity group are shown in Fig 6 (Left panel, a-d).

**Figure 6.**
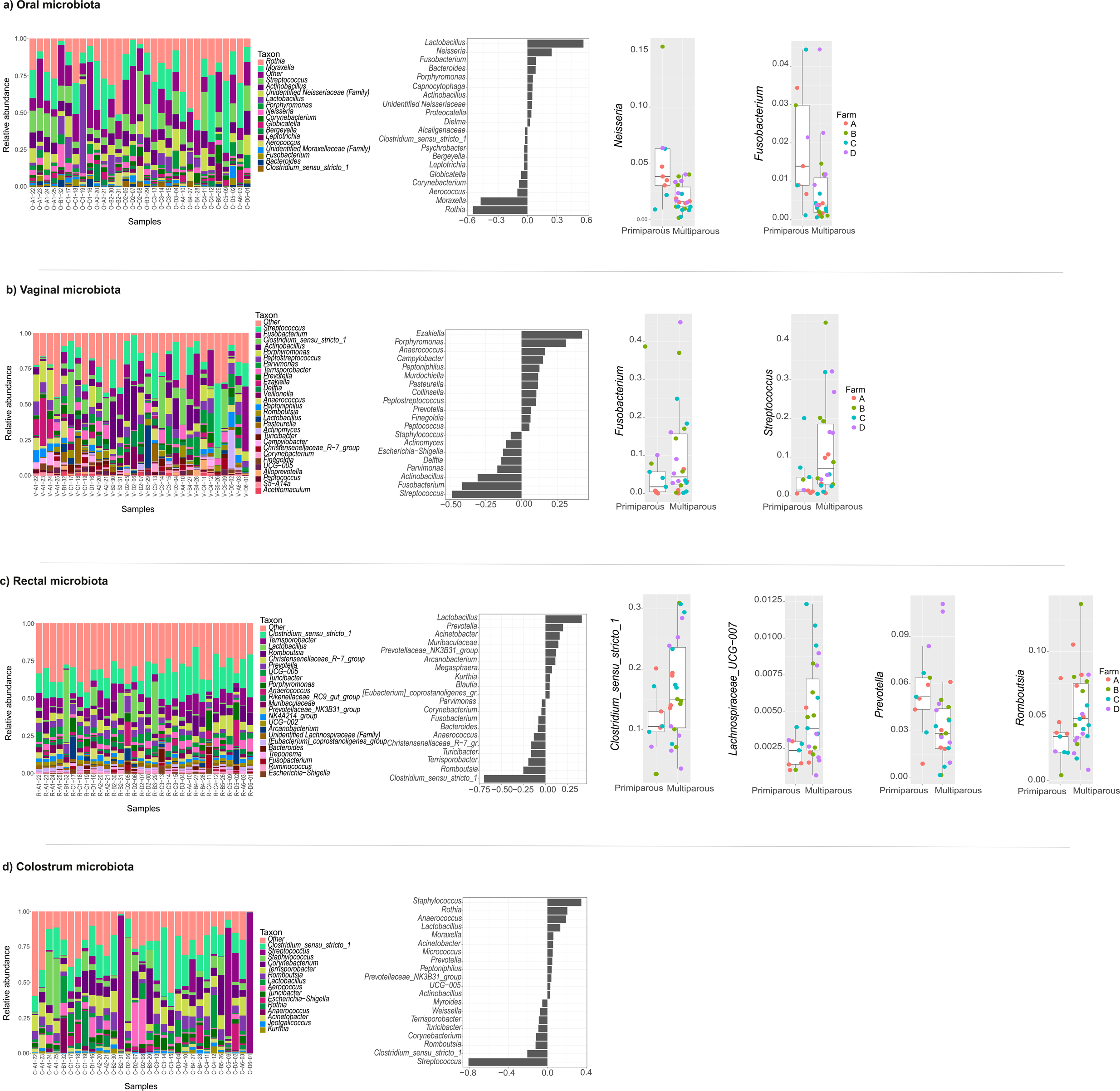
Differences in the microbial compositions between primiparous and multiparous sows from four Finnish commercial farms. a) oral, b) vaginal, c) rectal, d) colostrum samples. Left: Relative abundances of major genera, samples sorted by parity (A1-D1 = primiparous, A2-D6 = multiparous), middle: dbRDA analysis, right: Differences in relative abundances of specific genera between primiparous and multiparous sows.

Variation in the RAs of some of the major genera between individuals was extensive, especially of *Lactobacillus* in oral and rectal samples, *Fusobacterium* in vagina, and *Streptococcus* in colostrum. Distance-based redundancy analysis (dbRDA) shows the genera with largest contributions to the difference between the groups in each sample type (Fig 6, Middle panel). The genera with a tendency to higher RAs in multiparous sows point left and vice versa. The genera highlighted in dbRDA were tested for statistical significance and consistency of the difference across the farms. Fig 6 (Right panel) shows boxplots for the genera that were significantly and consistently different in each of the sample types.

In oral samples, the RAs of the genera *Fusobacterium* and *Neisseria* were significantly lower in multiparous compared with primiparous sows. The lower RA of the genus *Fusobacterium* in the oral samples of multiparous sows was due to ASVs likely representing several species. Among these, *F. necrophorum* and *F. nucleatum* were unequivocally identified based on BLAST sequence comparisons; the others were unknown. ASVs likely representing several species of *Neisseria* had lower RAs in multiparous sows. These were unknown or the identification was ambiguous.

In vaginal samples, there was a marked tendency towards higher RAs of the genera *Fusobacterium* and *Streptococcus* in multiparous compared with primiparous sows (Fig 6b Right panel). Around half of the samples in multiparous sows had higher RAs of these genera than primiparous sows. The RA of *Fusobacteria* was consistently higher in multiparous than in primiparous sows in the samples from all the four farms, although the difference was not statistically significant. The RAs of *Streptococcus* were higher in multiparous sows on three farms. The differences of the RAs of the other genera were either not significant between the groups or were not observed in all the farms. The differences in *Fusobacteria* were due to ASVs that were identified as *Fusobacterium gastrosuis* and *Fusobacterium necrophorum*. The RAs of several species of the genus *Streptococcus* differed between the groups.

In rectal samples, the RAs of the genera *Clostridium sensu stricto 1*, *Romboutsia*, and *Lachnospiraceae*_UCG_007 were significantly higher and *Prevotella* lower in multiparous compared with primiparous sows (Fig 6c Right panel). The difference was observed in samples from all the four farms. In contrast, the difference in the RA of *Lactobacillus* between the groups displayed in dbRDA was mainly due to the exceptionally high RA of *Lactobacillus* in only one primiparous sow. None of the ASVs with different levels in the two groups could be assigned to known species.

In colostrum, the RA of the genus *Streptococcus* was higher in multiparous than primiparous sows. This was due to colostrum samples from three multiparous sows having RAs of *Streptococcus* up to 75-98% (D6-01, B2-31 and C5-09 in Fig 6d Left panel). In addition, two samples (B4-27 and D5-02) had moderately high abundances of *Streptococcus* (20-22%). This was due to the high RA of an ASV matching the sequence of *Streptococcus dysgalactiae* in these samples. The average RA of this species in the other colostrum samples of multiparous sows was not higher than that of primiparous sows and remained < 0.2%. No clear relation in the abundances of *S. dysgalactiae* in colostrum versus vaginal samples for the affected individuals was detected. Sow D6-01 had higher than average RA (16%) of *S. dysgalactiae* also in the vagina, while the RAs for the other affected individuals were < 0.7%. ASVs matching to *S. dysgalactiae* were not detected in any of the rectal samples.

## Discussion

The aim of this study was to obtain an overview of the mucosal and colostrum microbiota of sows on Finnish commercial farms under normal production conditions. The other studies in this field are from settings that were different with respect to geographical location, climate, animal breeds, and animal husbandry. We characterized the microbial communities present on oral, vaginal, and rectal mucosa of late pregnant sows and in colostrum from four piglet-producing farms. We detected differences between the farms both in microbial diversity and the RAs of genera. We also observed differences between primiparous and multiparous sows in the RAs of various genera in all the sample types. Some of these differences were shared between all the farms, while others were limited to only some of them.

Moderate differences between the farms were detected both in microbial diversity and RAs of the main genera in all the sample types. Although standard Finnish animal husbandry procedures were used on all the farms, many factors can contribute to differences in microbiota compositions, such as the genetic background of the sows [31, 32], details in feeding, [25, 26, 33], antibiotic use and stress [27, 34], and characteristics of the physical and microbial environment [35], especially during early life when the microbiota of the sows was first established ( [24, 28, 36, 37].

### Oral microbiota

Proteobacteria was the most abundant phylum in the sow oral samples in contrast to the other sample types, which were dominated by Firmicutes. At the genus level, the oral microbiota was dominated by *Rothia*, *Moraxella*, *Streptococcus*, and *Actinobacillus*, followed by *Porphyromonas*, *Neisseria*, and *Corynebacterium*. Species belonging to these genera are commonly detected in the oral cavity and upper respiratory tract of various mammalian species, such as humans, dogs, and cattle [38-40]. Relatively few studies have addressed the composition of oral microbiota in pigs, mostly in piglets and growing pigs at various oral sites, such as tonsilla [41, 42], gingiva, buccal mucosa, and floor of the mouth [43]. The four top genera of our study were also identified as the most abundant in the saliva of sows and piglets in the study by Murase et al., where a similar procedure of sampling was used [44].

Overall, around half of the top 20 genera were shared between the two studies. The composition in our study also overlapped with two other studies on sow oral microbiota where cotton swabs [24] or ropes [45] were used in sample collection, with eight of our 20 top genera shared with those of each of the studies. *Acinetobacter* was the dominant genus in both of these studies, *Streptococcus* was among the most abundant genera, but *Rothia, Moraxella*, and *Actinobacillus* were either less abundant or not detected. Only the genera *Streptococcus*, *Lactobacillus* and *Clostridium sensu stricto 1* were among the 20 most abundant in all the four studies, indicating variability of the RAs and prevalences of the oral genera. The differences in the reported compositions may be due to many factors, such as feed and the environmental microbial exposure of the sows. Detailed information on the effect of the reproductive cycle on the sow oral microbiota composition is not yet available. Li et al. reported a transient increase of *Actinobacillus* during parturition [24]. Pregnancy is known to induce changes in the human oral microbiota, some of which may be related to adverse pregnancy outcomes [46].

We detected significantly higher abundances of the genera *Fusobacterium* and *Neisseria* in multiparous versus primiparous sows from the four farms. To the best of our knowledge, the effects of parity on the oral microbiota of sows have not been reported previously. The difference in *Fusobacterium* was attributed to *F. necrophorum*. *F. necrophorum* is present in the mouth, upper respiratory, and gastrointestinal tract and is an opportunistic pathogen in humans and other species ( [47, 48]. In pigs, *F. necrophorum* can cause necrotic stomatitis and facial necrosis [49, 50]. The species of *Neisseria* involved could not be determined without ambiguity. Several species of *Neisseria* are associated with periodontal health in humans [38] and dogs [39, 51].

### Vaginal microbiota

*Actinobacillus*, *Clostridium sensu stricto 1*, *Parvimonas*, and *Streptococcus* were among the most abundant genera in the vaginal microbiota. These genera have also been reported to belong to the most abundant in pregnant sows by others [27, 52, 53]. For the other top genera, there was more variation between the studies. *Actinobacillus* and *Clostridium* were also among the most abundant genera in weaning sows [54]. The RAs of Lactobacilli in our study were comparatively low but still within the range of RAs reported in the other studies. While lactobacilli are dominant in the human vaginal microbiota and have an important role in the maintenance of vaginal pH, the RAs of lactobacilli in other mammalian species, including pigs, are much lower [55, 56]).

A marked proportion of the multiparous sows had higher RAs of the genera *Fusobacterium* and *Streptococcus* than primiparous sows. Although not statistically significant at the group level, this difference was seen in all farms and in around half of the multiparous sows. The species corresponding to the differentially abundant ASVs were *F. gastrosuis* and *F. necrophorum*. *F. gastrosuis* is resident in the pig gastrointestinal tract and is implicated in gastric histopathology [57]. *F. necrophorum* is associated with metritis in dairy cows [58].

Parity-related differences in the sow vaginal microbiota have not been reported previously. Increased RAs of multiple genera have been reported in association with endometritis, including *Escherichia-Shigella*, *Bacteroides*, *Fusobacterium*, *Clostridium sensu stricto 1*, *Porphyromonas*, *Streptococcus*, *Actinobacillus*, *Burkholderia*, *Serratia*, and *Parvimonas* [53, 59, 60].

### Rectal microbiota

The major rectal genera in the studied sows were *Clostridium sensu stricto 1, Terrisporobacter*, *Lactobacilllus*, and *Romboutsia*. *Christensenellaceae*_R−7_group, *Prevotella*, *Turicibacter*, and *Rikenellaceae*_RC9_gut_group were also present in high relative abundances. These genera were also among the most abundant in other recent studies on sows from several geographically distinct locations and varying study designs [15-17, 19-21, 23, 27, 61, 62]. There is variation across studies in the genera reported as the most abundant in sow fecal or rectal samples, with *Clostridium* as the most abundant in some studies [19, 23, 62], including ours, and *Prevotella*, *Treponema*, or both in others [17, 20, 21]. However, all the genera defined as the minimal common fecal core in growing pigs by Holman et al. [63] (*Prevotella*, *Clostridium*, *Alloprevotella*, *Ruminococcus*, and the RC9 gut group) were among the most abundant both in our study and in the other sow studies. In addition to these, *Christensenellaceae*_R−7_group was among the most abundant in most of the sow studies.

Parity was associated with the RAs of several genera in rectal samples. Consistent differences included a significantly higher RA of *Clostridium sensu stricto 1*, *Romboutsia*, and *Lachnospiraceae*_UCG_007 and lower RA of *Prevotella* in multiparous compared with primiparous sows. Higher RA of *Clostridium sensu stricto 1* in third parity sows compared with first parity sows has also been reported earlier [23]. *Clostridium* species are commensal bacteria known as butyrate producers in the gut [64]. *Lachnospiraceae* are also producers of short-chain fatty acids [65]. *Prevotella* contains species with the ability to degrade plant glycans [66]. In growing pigs, *Prevotella* is a dominant genus that has multiple interactions with other microbial taxa; *Prevotella* species have effects on feeding efficiency [67]. Higher RAs of Prevotellaceae and Bacteroidota in high vs low parity have been observed previously [21]. Bacterial species belonging to the genus *Romboutsia* are gastrointestinal or environmental anaerobes identified from animal and human ileal samples [68, 69]. Little is known of their role in the gastrointestinal tract, but higher RAs of *Romboutsia* along with other commensal intestinal genera are associated with lower prevalence of type 2 diabetes and lower RAs with hypertension in humans [70, 71].

The differences in the abundance of the genus *Lactobacillus* detected in our study were due to only a few individuals and could not be considered as consistent over the data. Parity-related differences in the abundances of *Lactobacillus amylovorus* and *L. reuteri* have however been observed in pregnant sows by Berry et. al [22]. They also observed increased abundance of *Treponema bryantii* with higher parity. The genus *Treponema* was detected in our study, albeit with lower RA than in some other studies, and its RA did not differ between the parity groups. Several species of *Treponema* were present, but no ASVs corresponding to *T. bryantii* were detected.

### Colostrum microbiota

The most abundant genera in colostrum were *Clostridium sensu stricto 1*, *Streptococcus*, *Staphylococcus*, *Corynebacterium*, *Terrisporobacter*, *Romboutsia*, and *Lactobacillus*.

*Clostridium*, *Streptococcus*, *Staphylococcus*, and *Lactobacillus* were identified as the most abundant genera in other studies [28, 72]. *Pseudomonas* was reported as the core genus by Li et al., with variation between the study breeds for the abundances of other genera [24].

The main reason for the variation in colostrum samples between the parity groups was the high RA of the genus *Streptococcus* in five (of 23) multiparous animals from three different farms. This was due to the exceptionally high RA of the species *S. dysgalactiae*. This species is a known pathogen that causes mastitis in cows [73] and has been found in increased abundance in sows with purulent vaginal discharge [74]. When transmitted to newborn piglets it may cause arthritis and encephalitis [75]. The source of *S. dysgalactiae* in the colostrum of the study sows remains unknown. It was present in vaginal samples of both affected and nonaffected individuals but was not detected in rectal samples. Environmental transmission is also possible [73].

### Limitations of the study

This was an exploratory study on the compositions of mucosal and colostrum microbiota on sows reared under regular commercial production conditions. It was not specifically designed to study differences between sows of different parity. Therefore, the results presented must be considered preliminary. However, it is notable that our analysis uncovered differences between the rectal microbiotas of primiparous and multiparous sows that are similar to those previously reported from study settings specifically designed for that purpose.

## Conclusions

Our study presents the microbial compositions of the mucosal sites and colostrum of commercially reared pregnant sows from Finnish piglet-producing farms. Diverse microbiotas with some variations between farms were discovered, with *Rothia*, *Moraxella*, and *Streptococcus* as the major genera in oral, *Streptococcus* in vaginal, *Clostridium sensu stricto 1* in rectal, and *Streptococcus* in colostrum samples. Several potentially interesting differences between primiparous and multiparous sows were detected, involving *Fusobacterium* and *Neisseria* in oral, *Fusobacterium and Streptococcus* in vaginal, *Clostridium sensu stricto 1, Romboutsia*, *Lachnospiraceae_UCG_007*, *Prevotella* in rectal, and *Streptococcus* in colostrum samples. Further targeted studies are needed to define the differences related to parity and the impact of the sow microbiota on reproductive health.

## Materials and methods

### Farms, animals, and management

Four commercial piglet-producing farms in western and southwestern Finland participated voluntarily in the study during 2018-2019. The study farms are hereafter referred to as farms A-D.

Detailed information about expected farrowing days and sow parities of one farrowing group in each farm were obtained from the farmers. The first farm visit was planned to enable the researchers to supervise the maximum number of farrowings in three days on each farm. Otherwise, the farm staff followed their normal management practices during the study. The researchers inspected the study sows visually while supervising their farrowings. None of the sows were considered to have health concerns.

On each farm, the sows in the farrowing units were housed in farrowing pens (4.6-4.8 m^2^) with crates and partly slatted floors. No bedding material was used for the sows and piglets in the farrowing pens. Within the farrowing pen, the piglets had a separate nest with a heat lamp and solid floor. The sows were fed a standard liquid feed three times a day in farms A-C and in farm D three to four times before farrowing, twice right after farrowing, and four times daily during the rest of the lactation. Further details of the study farms, animals, and sow management are shown in Table 1.

**Table 1.**
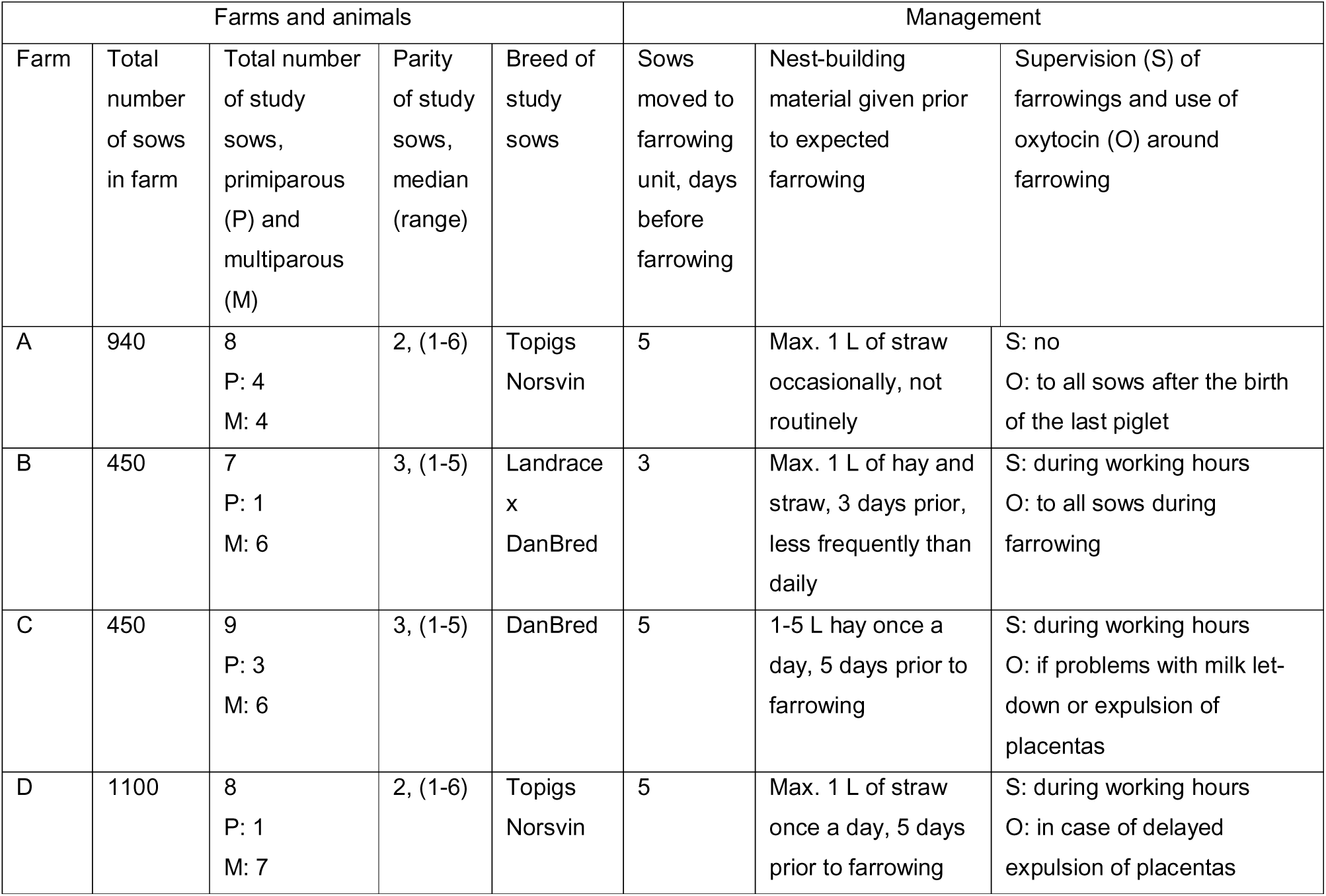
Descriptive information about the four study farms, animals, and management.

### Sampling

Swab samples from sow oral, vaginal, and rectal mucosa were collected from 32 pregnant sows on average 1.5 (SD 1.2) days prior to farrowing. Factory clean protective gloves were used during sampling. Sterile flocked swabs (FLOQSwabs®, Copan Diagnostics Inc, CA, USA) were used for oral and vaginal samples. Oral samples were taken by wiping the buccal mucosa, and vaginal samples by opening the vulva with one hand and wiping the mucosa just inside the vagina with the swab. From one sow the oral sample was not obtained. Rectal samples were taken with sterile cotton swabs (Applimed SA, Châtel-St-Denis). The swabs were placed into cryotubes that were placed in a cool box, moved to a –18°C freezer within an hour and further to a –80°C freezer on arrival at the laboratory. Colostrum samples were milked within 6 hours from birth of the first piglet into sterile 10-ml plastic tubes after disinfection of the udder as described in detail in [76]. From one sow the colostrum sample was not obtained. Colostrum samples were stored similarly as the other samples.

### DNA extraction

DNA from all sample types was extracted using a ZymoBIOMICS^TM^ DNA Miniprep Kit (Zymo Research, Irvine, CA) with minor modifications to the kit instructions as described below. Different sample types were processed at different times to avoid cross-contamination between samples.

All samples were thawed on ice on extraction day. The sampling swabs were transferred with sterile instruments into lysis tubes (ZR BashingBead Lysis Tubes, Zymo Research, Irvine, CA), and 750 µl of ZymoBiomics Lysis Solution was added to each tube. Bacterial cells were lysed three times with a Fastprep® 24 (5.5 m/s, 60 s); the tubes were then centrifuged 16000 x g for 5 min at 4°C. The supernatant was transferred into ultraclean Eppendorf tubes and 200 µl of lysis solution was added to the tubes. The fast-prep and centrifugation steps were repeated. The supernatants were combined and centrifuged at 8000 x g for 1 min at room temperature. Plain sterile flocked swabs (FLOQSwabs®, Copan Diagnostics Inc, CA, USA) were used as negative controls for vaginal swabs during three of four analysis rounds. For oral and rectal swabs, negative controls were not included in the extraction round.

For colostrum samples, 2 ml of thawed sample were transferred into ultraclean microcentrifuge tubes and centrifuged at 16000 x g for 10 min at +4 °C. Excess supernatant was removed until 200 µl of the supernatant remained. 750 µl of lysis solution (ZymoBiomics Lysis Solution) and 19 µl 20 mg/ml proteinase K solution in storage buffer (Zymo Research, Irvine, CA, USA) were added into each tube and mixed. The tubes were incubated at 55°C for 30 min. The remaining steps of the extraction from fast-prepping onward were performed as for the other sample types. Ultrapure water (Zymo Research, Irvine, CA, USA), stored in ultraclean microcentrifuge tubes in a similar way to the colostrum samples, was used as a negative control for colostrum samples.

After extraction, the DNA concentration of all sample types was measured with Qubit 3.0 Fluorometer (Thermo Fisher Scientific Inc., Waltham, MA).

### Library preparation and 16S rRNA gene amplicon sequencing

The hypervariable regions V3-V4 of the 16S ribosomal RNA (rRNA) genes were sequenced using the Illumina MiSeq platform in the DNA core facility of the University of Helsinki, essentially as described previously [77, 78]. Samples, negative controls, and ZymoBiomics Microbial Community Standard (Zymo Research, Irvine, CA, United States) were first amplified using 1× Phusion Hot Start II High-Fidelity PCR Master Mix (Thermo Scientific), 2.5% DMSO (Thermo Scientific), 500 nM of 16S rRNA V3 and V4 gene primers (341F and 785R; Metabion), and 1.25 μl of DNA extract. Total volume was 25 μl. The thermal cycling conditions included an initial denaturation step at 98°C for 30 s, 14-21 cycles of denaturing at 98°C for 10 s, annealing at 56°C for 30 s, and extension at 72°C for 20 s (14 cycles for fecal samples, 18 cycles for oral and vaginal samples, and 21 cycles for colostrum samples). The final extension step was at 72°C for 5 min. A T100™ Thermal Cycler (Bio-Rad Laboratories) was used. The amplicons were treated with exonuclease I and shrimp alkaline phosphatase for 30 min at 37°C to remove excess free primers. The second-round PCR amplifications were performed using an Illumina forward and reverse primer set, Phusion Hot-Start II polymerase, High Fidelity buffer, and 2.5% DMSO. The following thermal cycling conditions were applied with an Arktik thermal cycler (Finnzymes/Thermo Scientific): initial denaturation at 98°C for 30 s, 18 cycles at 98°C for 10 s, 65°C for 30 s, 72°C for 10s, and a final extension at 72°C for 5 min. Sample libraries were then pooled, and the pool was purified with a bead wash (MagSi-NGS Plus 0.9x). The final 16S rRNA gene amplicons were sequenced on an Illumina MiSeq sequencer using the v2 600 cycle kit paired-end (325 bp + 285 bp).

### Bioinformatics and statistics

The raw sequence data were processed as described in detail previously [78]. Briefly, the read quality was inspected using FastQC and MultiQC [79, 80]. Primers and spacers were trimmed using Cutadapt version 1.10 [81]. The mapping file was validated using Keemei [82]. QIIME2 version 2022.2 and the DADA2 plugin were used to de-noise and filter the reads, call amplicon sequence variants (ASVs), and generate a feature table [83, 84]. Taxonomy was assigned using the SILVA v138 99% database [85, 86]. Singleton sequences and sequences derived from chloroplasts or mitochondria were removed. One vaginal microbiota sample was excluded from further analyses as it was mostly composed of Delftia, a known reagent contaminant. Species-level identifications were confirmed by BLASTN 2.14.1+ search of the ASV sequences.

ASV data were analyzed using the R packages phyloseq 1.44.0 [87] mia 1.8.0 [88] and microbiome 1.23.1 [89]. Missing low-level taxonomic annotations (NA in phyloseq) were replaced with upper-level annotations using fantaxtic 2.0.1 [90]. Alpha diversity indices were calculated in phyloseq without rarefaction; rarefaction had no marked effect on the results. Statistical significances of alpha diversity differences were evaluated using the non-parametric Kruskal-Wallis rank-sum test and pairwise Wilcoxon rank-sum exact test with Bonferroni correction. Principal coordinates analyses (PCoA) and distance-based redundancy analysis (dbRDA) were performed using vegan 2.6-4 [91]. The core microbiota heatmaps were generated using the microbiome package. Scater 1.18.6 package [92] was used to generate the violin plots. Color-blind friendly palettes were obtained from *I Want Hue* [93]. Species-level identifications were confirmed by a search of the corresponding ASV sequences against nucleotide sequence database using BLASTN 2.14.1+. Only identifications with 100% match with the specific species and no matches with > 98% similarity to any other species were considered valid.

### Ethics approval

The experiment was conducted under the EU legislation on the protection of animals used for scientific purposes (Directive 2010/63/EU) and approved by the Southern Finland Regional State Administrative Agency (ESAVI/16950/2018). The farm owners provided signed written informed consent at the beginning of the study.

## Supporting information

Table S1. Overview of the sequencing data.

## Availability of data

The raw 16S rRNA gene amplicon sequencing dataset has been deposited in the NCBI Sequence Read Archive at https://www.ncbi.nlm.nih.gov/sra/PRJNA1069904 with the accession number PRJNA1069904.

## Author’s contributions

All authors contributed to planning the study. VP, EK, MH, and MN collected the samples. VP and EK performed the DNA extraction and AH and MN the bioinformatic processing of the data. MN and AH performed the statistical processing of the data. MN and TPM analyzed and compiled the results, EK, MN and TPM visualized the data. TPM and VP wrote the manuscript with contributions from all authors. All authors read and commented on the manuscript and approved the final version.

## Acknowledgements

The authors wish to thank Kirsi Lahti for excellent technical assistance and the University of Helsinki HiLife DNA sequencing and genomics core facility for performing the 16S RNA gene amplicon sequencing.

## References

1. Chu DM, Ma J, Prince AL, Antony KM, Seferovic MD, Aagaard KM. Maturation of the infant microbiome community structure and function across multiple body sites and in relation to mode of delivery. Nat Med. 2017;23(3):314–26; doi: 10.1038/nm.4272.

2. Gilbert JA, Blaser MJ, Caporaso JG, Jansson JK, Lynch SV, Knight R. Current understanding of the human microbiome. Nat Med. 2018;24(4):392–400; doi: 10.1038/nm.4517.

3. Zaidi S, Ali K, Khan AU. It’s all relative: analyzing microbiome compositions, its significance, pathogenesis and microbiota derived biofilms: Challenges and opportunities for disease intervention. Arch Microbiol. 2023;205(7):257; doi: 10.1007/s00203-023-03589-7.

4. Kayama H, Okumura R, Takeda K. Interaction Between the Microbiota, Epithelia, and Immune Cells in the Intestine. Annu Rev Immunol. 2020;38:23–48; doi: 10.1146/annurev-immunol-070119-115104.

5. Pickard JM, Zeng MY, Caruso R, Nunez G. Gut microbiota: Role in pathogen colonization, immune responses, and inflammatory disease. Immunol Rev. 2017;279(1):70–89; doi: 10.1111/imr.12567.

6. Stokes CR. The development and role of microbial-host interactions in gut mucosal immune development. J Anim Sci Biotechnol. 2017;8:12; doi: 10.1186/s40104-016-0138-0.

7. Belkaid Y, Harrison OJ. Homeostatic Immunity and the Microbiota. Immunity. 2017;46(4):562–76; doi: 10.1016/j.immuni.2017.04.008.

8. Zheng D, Liwinski T, Elinav E. Interaction between microbiota and immunity in health and disease. Cell Res. 2020;30(6):492–506; doi: 10.1038/s41422-020-0332-7.

9. Graham DB, Xavier RJ. Conditioning of the immune system by the microbiome. Trends Immunol. 2023;44(7):499–511; doi: 10.1016/j.it.2023.05.002.

10. Wang H, Xu R, Zhang H, Su Y, Zhu W. Swine gut microbiota and its interaction with host nutrient metabolism. Anim Nutr. 2020;6(4):410–20; doi: 10.1016/j.aninu.2020.10.002.

11. Morais LH, Schreiber HLt, Mazmanian SK. The gut microbiota-brain axis in behaviour and brain disorders. Nat Rev Microbiol. 2021;19(4):241–55; doi: 10.1038/s41579-020-00460-0.

12. Verbeek E, Keeling L, Landberg R, Lindberg JE, Dicksved J. The gut microbiota and microbial metabolites are associated with tail biting in pigs. Sci Rep. 2021;11(1):20547; doi: 10.1038/s41598-021-99741-8.

13. Patil Y, Gooneratne R, Ju XH. Interactions between host and gut microbiota in domestic pigs: a review. Gut Microbes. 2020;11(3):310–34; doi: 10.1080/19490976.2019.1690363.

14. Luo Y RW, Smidt H, Wright AG, Yu B, Schyns G, McCormack UM, Cowieson AJ, Yu J, He J, Yan H, Wu J, Mackie RI, Chen D.. Dynamic Distribution of Gut Microbiota in Pigs at Different Growth Stages: Composition and Contribution. Microbiol Spectr. 2022;10(3); doi: 10.1128/spectrum.00688-21.

15. Zhou P, Zhao Y, Zhang P, Li Y, Gui T, Wang J, et al. Microbial Mechanistic Insight into the Role of Inulin in Improving Maternal Health in a Pregnant Sow Model. Frontiers in Microbiology. 2017;8; doi: 10.3389/fmicb.2017.02242.

16. Cheng C, Wei H, Yu H, Xu C, Jiang S, Peng J. Metabolic Syndrome During Perinatal Period in Sows and the Link With Gut Microbiota and Metabolites. Front Microbiol. 2018;9:1989; doi: 10.3389/fmicb.2018.01989.

17. Ji YJ, Li H, Xie PF, Li ZH, Li HW, Yin YL, et al. Stages of pregnancy and weaning influence the gut microbiota diversity and function in sows. J Appl Microbiol. 2019;127(3):867–79; doi: 10.1111/jam.14344.

18. Huang X, Gao J, Zhao Y, He M, Ke S, Wu J, et al. Dramatic Remodeling of the Gut Microbiome Around Parturition and Its Relationship With Host Serum Metabolic Changes in Sows. Front Microbiol. 2019;10:2123; doi: 10.3389/fmicb.2019.02123.

19. Sun L, Zhang Y, Chen W, Lan T, Wang Y, Wu Y, et al. The Dynamic Changes of Gut Microbiota during the Perinatal Period in Sows. Animals (Basel). 2020;10(12); doi: 10.3390/ani10122254.

20. Fu H, He M, Wu J, Zhou Y, Ke S, Chen Z, et al. Deep Investigating the Changes of Gut Microbiome and Its Correlation With the Shifts of Host Serum Metabolome Around Parturition in Sows. Front Microbiol. 2021;12:729039; doi: 10.3389/fmicb.2021.729039.

21. Gaukroger CH, Edwards SA, Walshaw J, Nelson A, Adams IP, Stewart CJ, et al. Shifting sows: longitudinal changes in the periparturient faecal microbiota of primiparous and multiparous sows. Animal. 2021;15(3):100135; doi: 10.1016/j.animal.2020.100135.

22. Berry ASF, Pierdon MK, Misic AM, Sullivan MC, O’Brien K, Chen Y, et al. Remodeling of the maternal gut microbiome during pregnancy is shaped by parity. Microbiome. 2021;9(1):146; doi: 10.1186/s40168-021-01089-8.

23. Greiner LL, Humphrey DC, Holland SN, Anderson CJ, Schmitz-Esser S. The validation of the existence of the entero-mammary pathway and the assessment of the differences of the pathway between first and third parity sows. Transl Anim Sci. 2022;6(2):txac047; doi: 10.1093/tas/txac047.

24. Li Y, Liu Y, Ma Y, Ge X, Zhang X, Cai C, et al. Effects of Maternal Factors and Postpartum Environment on Early Colonization of Intestinal Microbiota in Piglets. Front Vet Sci. 2022;9:815944; doi: 10.3389/fvets.2022.815944.

25. Hasan S, Junnikkala S, Peltoniemi O, Paulin L, Lyyski A, Vuorenmaa J, et al. Dietary supplementation with yeast hydrolysate in pregnancy influences colostrum yield and gut microbiota of sows and piglets after birth. PLoS One. 2018;13(5):e0197586; doi: 10.1371/journal.pone.0197586.

26. Liu B ZX, Cui Y, Wang W, Liu H, Li, Z GZ, Ma S, Li D, Wang C, Shi Y. Consumption of dietary fiber from different sources during pregnancy alters sow gut microbiota and improves performance and reduces inflammation in sows and piglets. mSystems. 2021;6(1); doi: 10.1128/mSystems.00591-20.

27. He J, Zheng W, Tao C, Guo H, Xue Y, Zhao R, et al. Heat stress during late gestation disrupts maternal microbial transmission with altered offspring’s gut microbial colonization and serum metabolites in a pig model. Environ Pollut. 2020;266(Pt 3):115111; doi: 10.1016/j.envpol.2020.115111.

28. Chen X, Xu J, Ren E, Su Y, Zhu W. Co-occurrence of early gut colonization in neonatal piglets with microbiota in the maternal and surrounding delivery environments. Anaerobe. 2018;49:30–40; doi: 10.1016/j.anaerobe.2017.12.002.

29. Chen W, Ma J, Jiang Y, Deng L, Lv N, Gao J, et al. Selective Maternal Seeding and Rearing Environment From Birth to Weaning Shape the Developing Piglet Gut Microbiome. Front Microbiol. 2022;13:795101; doi: 10.3389/fmicb.2022.795101.

30. Guevarra RB, Lee JH, Lee SH, Seok MJ, Kim DW, Kang BN, et al. Piglet gut microbial shifts early in life: causes and effects. J Anim Sci Biotechnol. 2019;10:1; doi: 10.1186/s40104-018-0308-3.

31. Pajarillo EA, Chae JP, Balolong MP, Kim HB, Seo KS, Kang DK. Pyrosequencing-based analysis of fecal microbial communities in three purebred pig lines. J Microbiol. 2014;52(8):646–51; doi: 10.1007/s12275-014-4270-2.

32. Yang H, Xiao Y, Wang J, Xiang Y, Gong Y, Wen X, et al. Core gut microbiota in Jinhua pigs and its correlation with strain, farm and weaning age. J Microbiol. 2018;56(5):346–55; doi: 10.1007/s12275-018-7486-8.

33. Tilocca B, Burbach K, Heyer CME, Hoelzle LE, Mosenthin R, Stefanski V, et al. Dietary changes in nutritional studies shape the structural and functional composition of the pigs’ fecal microbiome-from days to weeks. Microbiome. 2017;5(1):144; doi: 10.1186/s40168-017-0362-7.

34. Schokker D, Zhang J, Zhang LL, Vastenhouw SA, Heilig HG, Smidt H, et al. Early-life environmental variation affects intestinal microbiota and immune development in new-born piglets. PLoS One. 2014;9(6):e100040; doi: 10.1371/journal.pone.0100040.

35. Megahed A, Zeineldin M, Evans K, Maradiaga N, Blair B, Aldridge B, et al. Impacts of environmental complexity on respiratory and gut microbiome community structure and diversity in growing pigs. Sci Rep. 2019;9(1):13773; doi: 10.1038/s41598-019-50187-z.

36. Nowland TL, Plush KJ, Barton M, Kirkwood RN. Development and Function of the Intestinal Microbiome and Potential Implications for Pig Production. Animals (Basel). 2019;9(3); doi: 10.3390/ani9030076.

37. Luhrmann A, Ovadenko K, Hellmich J, Sudendey C, Belik V, Zentek J, et al. Characterization of the fecal microbiota of sows and their offspring from German commercial pig farms. PLoS One. 2021;16(8):e0256112; doi: 10.1371/journal.pone.0256112.

38. Yamashita Y, Takeshita T. The oral microbiome and human health. Journal of Oral Science. 2017;59(2):201–6; doi: 10.2334/josnusd.16-0856.

39. Wallis C, Marshall M, Colyer A, O’Flynn C, Deusch O, Harris S. A longitudinal assessment of changes in bacterial community composition associated with the development of periodontal disease in dogs. Vet Microbiol. 2015;181(3-4):271–82; doi: 10.1016/j.vetmic.2015.09.003.

40. McMullen C, Alexander TW, Leguillette R, Workentine M, Timsit E. Topography of the respiratory tract bacterial microbiota in cattle. Microbiome. 2020;8(1):91; doi: 10.1186/s40168-020-00869-y.

41. Lowe BA, Marsh TL, Isaacs-Cosgrove N, Kirkwood RN, Kiupel M, Mulks MH. Defining the “core microbiome” of the microbial communities in the tonsils of healthy pigs. BMC Microbiol. 2012;12:20; doi: 10.1186/1471-2180-12-20.

42. Pena Cortes LC, LeVeque RM, Funk J, Marsh TL, Mulks MH. Development of the tonsillar microbiome in pigs from newborn through weaning. BMC Microbiol. 2018;18(1):35; doi: 10.1186/s12866-018-1176-x.

43. Jeon JH, Lourenco JM, Fagan MM, Welch CB, Sneed SE, Dubrof S, et al. Changes in Oral Microbial Diversity in a Piglet Model of Traumatic Brain Injury. Brain Sci. 2022;12(8); doi: 10.3390/brainsci12081111.

44. Murase K, Watanabe T, Arai S, Kim H, Tohya M, Ishida-Kuroki K, et al. Characterization of pig saliva as the major natural habitat of Streptococcus suis by analyzing oral, fecal, vaginal, and environmental microbiota. PLoS One. 2019;14(4):e0215983; doi: 10.1371/journal.pone.0215983.

45. Hattab J, Marruchella G, Pallavicini A, Gionechetti F, Mosca F, Trachtman AR, et al. Insights into the Oral Bacterial Microbiota of Sows. Microorganisms. 2021;9(11); doi: 10.3390/microorganisms9112314.

46. Ye C, Kapila Y. Oral microbiome shifts during pregnancy and adverse pregnancy outcomes: Hormonal and Immunologic changes at play. Periodontol 2000. 2021;87(1):276-81; doi: 10.1111/prd.12386.

47. Lee WS, Jean SS, Chen FL, Hsieh SM, Hsueh PR. Lemierre’s syndrome: A forgotten and re-emerging infection. J Microbiol Immunol Infect. 2020;53(4):513–7; doi: 10.1016/j.jmii.2020.03.027.

48. Nagaraja TG, Narayanan SK, Stewart GC, Chengappa MM. Fusobacterium necrophorum infections in animals: pathogenesis and pathogenic mechanisms. Anaerobe. 2005;11(4):239–46; doi: 10.1016/j.anaerobe.2005.01.007.

49. Thomson JR, Friendship RM. Digestive System. In: Zimmerman JJ KL, Ramirez A, Schwartz, Stevenson GW, Zhang J., editor. Diseases of Swine. Hoboken NJ: John Wiley & Sons, Inc.; 2019. p. 234–63.

50. Torrison J, Cameron R. Integumentary System. In: Zimmerman JJ KL, Ramirez A, Schwartz, Stevenson GW, Zhang J., editor. Diseases of Swine. Hoboken NJ: John Wiley & Sons, Inc.; 2019. p. 292–312.

51. Tavares MO, Dos Reis LD, Lopes WR, Schwarz LV, Rocha RKM, Scariot FJ, et al. Bacterial community associated with gingivitis and periodontitis in dogs. Res Vet Sci. 2023;162:104962; doi: 10.1016/j.rvsc.2023.104962.

52. Torres Luque A, Fontana C, Pasteris SE, Bassi D, Cocconcelli PS, Otero MC. Vaginal bacterial diversity from healthy gilts and pregnant sows subjected to natural mating or artificial insemination. Res Vet Sci. 2021;140:26–37; doi: 10.1016/j.rvsc.2021.07.023.

53. Liang H, Cai R, Li C, Glendon OHM, Chengcheng H, Yan H. High-throughput sequencing of 16S rRNA gene analysis reveals novel taxonomic diversity among vaginal microbiota in healthy and affected sows with endometritis. Res Vet Sci. 2022;143:33–40; doi: 10.1016/j.rvsc.2021.12.003.

54. Zhang J, Liu M, Ke S, Huang X, Fang S, He M, et al. Gut and Vagina Microbiota Associated With Estrus Return of Weaning Sows and Its Correlation With the Changes in Serum Metabolites. Front Microbiol. 2021;12:690091; doi: 10.3389/fmicb.2021.690091.

55. Lorenzen E, Kudirkiene E, Gutman N, Grossi AB, Agerholm JS, Erneholm K, et al. The vaginal microbiome is stable in prepubertal and sexually mature Ellegaard Gottingen Minipigs throughout an estrous cycle. Vet Res. 2015;46:125; doi: 10.1186/s13567-015-0274-0.

56. Miller EA, Beasley DE, Dunn RR, Archie EA. Lactobacilli Dominance and Vaginal pH: Why Is the Human Vaginal Microbiome Unique? Front Microbiol. 2016;7:1936; doi: 10.3389/fmicb.2016.01936.

57. De Witte C, Demeyere K, De Bruyckere S, Taminiau B, Daube G, Ducatelle R, et al. Characterization of the non-glandular gastric region microbiota in Helicobacter suis-infected versus non-infected pigs identifies a potential role for Fusobacterium gastrosuis in gastric ulceration. Vet Res. 2019;50(1):39; doi: 10.1186/s13567-019-0656-9.

58. Cunha F, Jeon SJ, Daetz R, Vieira-Neto A, Laporta J, Jeong KC, et al. Quantifying known and emerging uterine pathogens, and evaluating their association with metritis and fever in dairy cows. Theriogenology. 2018;114:25–33; doi: 10.1016/j.theriogenology.2018.03.016.

59. Wang J, Li C, Nesengani LT, Gong Y, Zhang S, Lu W. Characterization of vaginal microbiota of endometritis and healthy sows using high-throughput pyrosequencing of 16S rRNA gene. Microb Pathog. 2017;111:325–30; doi: 10.1016/j.micpath.2017.08.030.

60. Zhang L, Wang L, Dai Y, Tao T, Wang J, Wu Y, et al. Effect of Sow Intestinal Flora on the Formation of Endometritis. Front Vet Sci. 2021;8:663956; doi: 10.3389/fvets.2021.663956.

61. Shao Y, Zhou J, Xiong X, Zou L, Kong X, Tan B, et al. Differences in Gut Microbial and Serum Biochemical Indices Between Sows With Different Productive Capacities During Perinatal Period. Front Microbiol. 2019;10:3047; doi: 10.3389/fmicb.2019.03047.

62. Uryu H, Tsukahara T, Ishikawa H, Oi M, Otake S, Yamane I, et al. Comparison of Productivity and Fecal Microbiotas of Sows in Commercial Farms. Microorganisms. 2020;8(10); doi: 10.3390/microorganisms8101469.

63. Holman DB, Brunelle BW, Trachsel J, Allen HK. Meta-analysis To Define a Core Microbiota in the Swine Gut. mSystems. 2017;2(3); doi: 10.1128/mSystems.00004-17.

64. Guo P, Zhang K, Ma X, He P. Clostridium species as probiotics: potentials and challenges. J Anim Sci Biotechnol. 2020;11:24; doi: 10.1186/s40104-019-0402-1.

65. Abdugheni R, Wang WZ, Wang YJ, Du MX, Liu FL, Zhou N, et al. Metabolite profiling of human-originated Lachnospiraceae at the strain level. iMeta. 2022;1(4); doi: 10.1002/imt2.58.

66. Ley RE. Gut microbiota in 2015: Prevotella in the gut: choose carefully. Nat Rev Gastroenterol Hepatol. 2016;13(2):69-70; doi: 10.1038/nrgastro.2016.4.

67. Amat S, Lantz H, Munyaka PM, Willing BP. Prevotella in Pigs: The Positive and Negative Associations with Production and Health. Microorganisms. 2020;8(10); doi: 10.3390/microorganisms8101584.

68. Gerritsen J, Fuentes S, Grievink W, van Niftrik L, Tindall BJ, Timmerman HM, et al. Characterization of Romboutsia ilealis gen. nov., sp. nov., isolated from the gastro-intestinal tract of a rat, and proposal for the reclassification of five closely related members of the genus Clostridium into the genera Romboutsia gen. nov., Intestinibacter gen. nov., Terrisporobacter gen. nov. and Asaccharospora gen. nov. Int J Syst Evol Microbiol. 2014;64(Pt 5):1600–16; doi: 10.1099/ijs.0.059543-0.

69. Gerritsen J, Umanets A, Staneva I, Hornung B, Ritari J, Paulin L, et al. Romboutsia hominis sp. nov., the first human gut-derived representative of the genus Romboutsia, isolated from ileostoma effluent. Int J Syst Evol Microbiol. 2018;68(11):3479–86; doi: 10.1099/ijsem.0.003012.

70. Chen Z, Radjabzadeh D, Chen L, Kurilshikov A, Kavousi M, Ahmadizar F, et al. Association of Insulin Resistance and Type 2 Diabetes With Gut Microbial Diversity: A Microbiome-Wide Analysis From Population Studies. JAMA Netw Open. 2021;4(7):e2118811; doi: 10.1001/jamanetworkopen.2021.18811.

71. Naik SS, Ramphall S, Rijal S, Prakash V, Ekladios H, Mulayamkuzhiyil Saju J, et al. Association of Gut Microbial Dysbiosis and Hypertension: A Systematic Review. Cureus. 2022;14(10):e29927; doi: 10.7759/cureus.29927.

72. Llaurado-Calero E, Climent E, Chenoll E, Ballester M, Badiola I, Lizardo R, et al. Influence of dietary n-3 long-chain fatty acids on microbial diversity and composition of sows’ feces, colostrum, milk, and suckling piglets’ feces. Front Microbiol. 2022;13:982712; doi: 10.3389/fmicb.2022.982712.

73. Wente N, Krömker V. Streptococcus dysgalactiae—Contagious or Environmental? Animals. 2020;10(11); doi: 10.3390/ani10112185.

74. Poor AP, Moreno LZ, Monteiro MS, Matajira CEC, Dutra MC, Leal DF, et al. Vaginal microbiota signatures in healthy and purulent vulvar discharge sows. Scientific Reports. 2022;12(1); doi: 10.1038/s41598-022-13090-8.

75. Oh S-I, Kim JW, Jung J-Y, Chae M, Lee Y-R, Kim JH, et al. Pathologic and molecular characterization ofStreptococcus dysgalactiaesubsp.equisimilisinfection in neonatal piglets. Journal of Veterinary Science. 2018;19(2); doi: 10.4142/jvs.2018.19.2.313.

76. Konig E, Sali V, Heponiemi P, Salminen S, Valros A, Junnikkala S, et al. Herd-Level and Individual Differences in Fecal Lactobacilli Dynamics of Growing Pigs. Animals (Basel). 2021;11(1); doi: 10.3390/ani11010113.

77. Alipour MJ, Jalanka J, Pessa-Morikawa T, Kokkonen T, Satokari R, Hynonen U, et al. The composition of the perinatal intestinal microbiota in cattle. Sci Rep. 2018;8(1):10437; doi: 10.1038/s41598-018-28733-y.

78. Husso A, Jalanka J, Alipour MJ, Huhti P, Kareskoski M, Pessa-Morikawa T, et al. The composition of the perinatal intestinal microbiota in horse. Sci Rep. 2020;10(1):441; doi: 10.1038/s41598-019-57003-8.

79. Andrews S, Krueger, F., Seconds-Pichon, A., Biggins, F., Wingett, S., : FastQC. A Quality Control Tool for High Throughput Sequence Data. In.; 2015.

80. Ewels P, Magnusson M, Lundin S, Kaller M. MultiQC: summarize analysis results for multiple tools and samples in a single report. Bioinformatics. 2016;32(19):3047–8; doi: 10.1093/bioinformatics/btw354.

81. Martin M. Cutadapt removes adapter sequences from high-throughput sequencing reads. EMBnetjournal. 2011;17(n. 1):10–2; doi: 10.14806/ej.17.1.200.

82. Rideout JR, Chase JH, Bolyen E, Ackermann G, Gonzalez A, Knight R, et al. Keemei: cloud-based validation of tabular bioinformatics file formats in Google Sheets. Gigascience. 2016;5:27; doi: 10.1186/s13742-016-0133-6.

83. Callahan BJ, McMurdie PJ, Rosen MJ, Han AW, Johnson AJ, Holmes SP. DADA2: High-resolution sample inference from Illumina amplicon data. Nat Methods. 2016;13(7):581–3; doi: 10.1038/nmeth.3869.

84. Bolyen E, Rideout JR, Dillon MR, Bokulich NA, Abnet CC, Al-Ghalith GA, et al. Reproducible, interactive, scalable and extensible microbiome data science using QIIME 2. Nat Biotechnol. 2019;37(8):852–7; doi: 10.1038/s41587-019-0209-9.

85. Quast C, Pruesse E, Yilmaz P, Gerken J, Schweer T, Yarza P, et al. The SILVA ribosomal RNA gene database project: improved data processing and web-based tools. Nucleic Acids Research. 2012;41(D1):D590–D6; doi: 10.1093/nar/gks1219.

86. Bokulich NA, Kaehler BD, Rideout JR, Dillon M, Bolyen E, Knight R, et al. Optimizing taxonomic classification of marker-gene amplicon sequences with QIIME 2’s q2-feature-classifier plugin. Microbiome. 2018;6(1):90; doi: 10.1186/s40168-018-0470-z.

87. McMurdie PJ, Holmes S. phyloseq: an R package for reproducible interactive analysis and graphics of microbiome census data. PLoS One. 2013;8(4):e61217; doi: 10.1371/journal.pone.0061217.

88. Ernst F SS, Borman T, Lahti L Mia: Microbiome analysis. R package version 1.8.0. 2023. Available from https://github.com/microbiome/mia

89. Lahti L, Shetty, S.: Tools for microbiome analysis in R. Microbiome package version 1.23.1. Bioconductor; 2017. Available from https://github.com/microbiome/microbiome

90. Teunisse GM: Fantaxtic - Nested Bar Plots for Phyloseq Data (Version 2.0.1) 2022. Available from https://github.com/gmteunisse/Fantaxtic

91. Oksanen J SG, Blanchet F, Kindt R, Legendre P, Minchin P, O’Hara R, Solymos P, Stevens M, Szoecs E, Wagner H, Barbour M, Bedward M, Bolker B, Borcard D, Carvalho G, Chirico M, De Caceres M, Durand S, Evangelista H, FitzJohn R, Friendly M, Furneaux B, Hannigan G, Hill M, Lahti L, McGlinn D, Ouellette M, Ribeiro Cunha E, Smith T, Stier A, Ter Braak C, Weedon J: _vegan: Community Ecology Package_. R package version 2.6–4. 2022. Available from https://CRAN.R-project.org/package=vegan

92. McCarthy DJ, Campbell KR, Lun AT, Wills QF. Scater: pre-processing, quality control, normalization and visualization of single-cell RNA-seq data in R. Bioinformatics. 2017;33(8):1179–86; doi: 10.1093/bioinformatics/btw777.

93. Mathieu J. I want hue. 2023. Available from https://github.com/medialab/iwanthue

