## Supplementary material for "Bacterial profiles of the oral, vaginal, and rectal mucosa and colostrum of periparturient sows": Table S1. Overview of the sequencing data.

**Additional file 1**

**Table S1.** Overview of the sequencing data.

| Sample type | n | Nr of reads (Average ± SD) | Asv (Average ± SD) |
|---|---|---|---|
| Oral | 31 | 49570 ± 9217 | 339 ± 71 |
| Vaginal | 31 | 61513 ± 9687 | 351 ± 221 |
| Rectal | 32 | 44881 ± 6380 | 625 ± 172 |
| Colostrum | 31 | 58722 ± 15543 | 404 ± 188 |
