## Supplementary figure 1. for "Bacterial profiles of the oral, vaginal, and rectal mucosa and colostrum of periparturient sows"

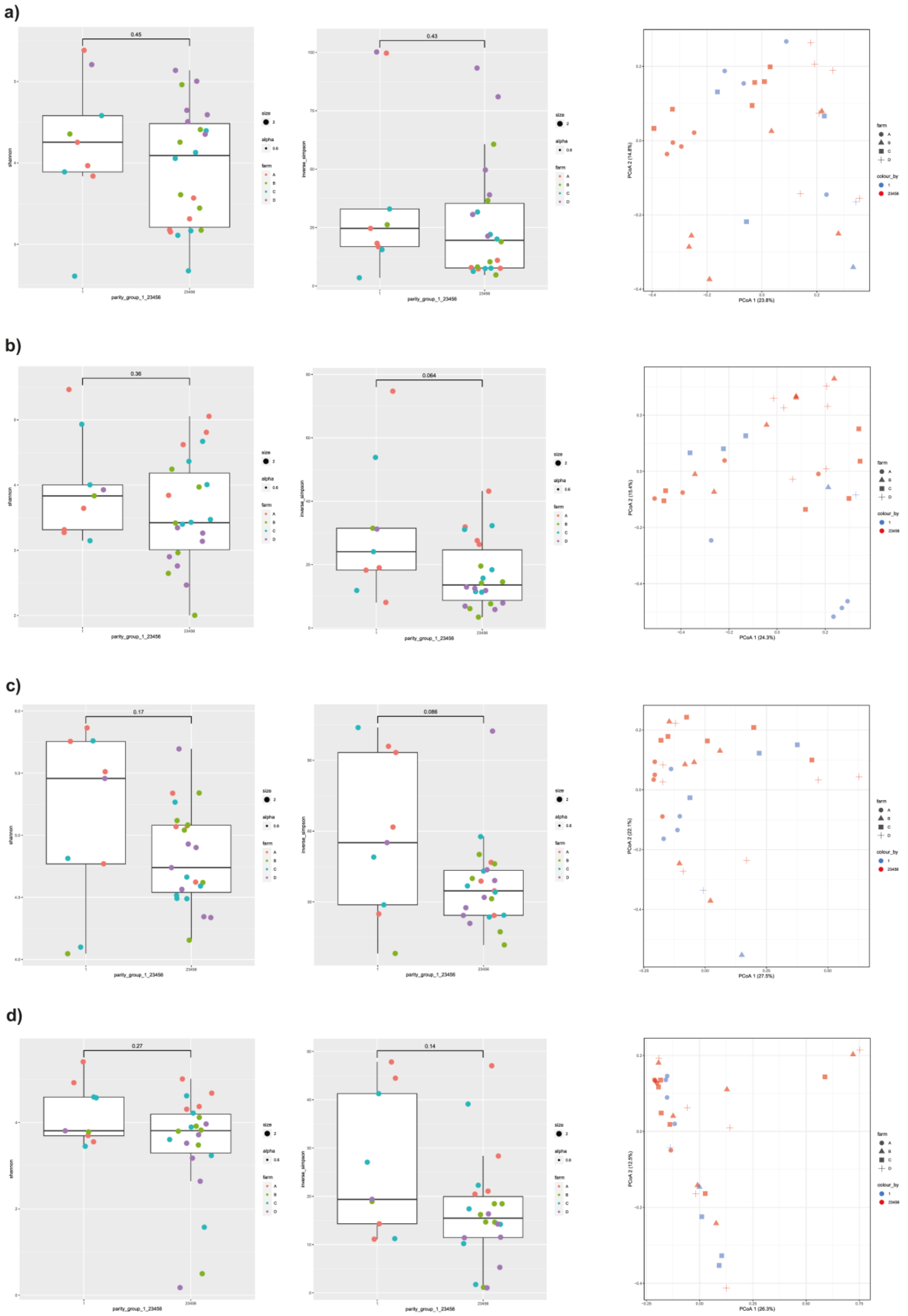


**Figure S1.** Comparison of the alpha and beta diversities of a) oral, b) vaginal, c) rectal and d) colostrum microbiota of primiparous vs multiparous sows. Alpha diversity index Shannon left, Inverse Simpson middle, and principal coordinates analysis based on Bray-Curtis dissimilarities left.
